# Electroconvulsive seizures for alcohol use disorder: a preclinical study

**DOI:** 10.64898/2026.03.30.715248

**Authors:** Rubén García-Cabrerizo, Pedro Bergas-Cladera, Carles Colom-Rocha, M. Julia García-Fuster

**Affiliations:** IUNICS, University of the Balearic Islands, Palma, Spain; Health Research Institute of the Balearic Islands (IdISBa), Palma, Spain; Department of Medicine, University of the Balearic Islands, Palma, Spain

**Keywords:** Addiction risk factors, early drug use, adolescence, alcohol use disorder, therapeutic options, neuromodulators

## Abstract

The use of neuromodulation techniques for the treatment of alcohol use disorder is receiving increasing attention, especially non-invasive approaches, such as repetitive transcranial magnetic stimulation or transcranial direct current stimulation, while the hypothetical use of electroconvulsive therapy remains unexplored. Given our experience inducing electroconvulsive seizures (ECS) for therapeutic purposes in psychopathology rodent models, we evaluated the role of ECS on reducing the increased voluntary ethanol consumption caused by adolescent ethanol exposure in our validated preclinical model. Rats were treated in adolescence with a binge paradigm of ethanol (2 g/kg, i.p.; 3 rounds of 2 days at 48-h intervals; post-natal day, PND 29-30, PND 33-34 and PND 37-38) or saline. Following persistent withdrawal until adulthood, rats were allowed to: voluntarily drink ethanol (20%) by a two-bottle choice test, for 3 days (PND 80-82); treated with ECS (95 mA for 0.6 s, 100 Hz, pulse width 0.6 ms; ear-clip electrodes) or SHAM for 5 days (PND 86-90); re-exposed to voluntarily ethanol exposure (PND 94-96). Brains were collected on PND 97 to evaluate hippocampal markers of ethanol toxicity and/or treatment response (e.g., NeuroD, NF-L, BDNF and NF-L/BDNF ratio). Our results reproduced the increased voluntary ethanol consumption in adult rats induced by adolescent ethanol exposure and demonstrated that ECS could improve this abuse-prone response. Moreover, we suggested a possible role for BDNF in the beneficial effects induced by ECS, especially reducing the neurotoxic ratio NF-L/BDNF. Overall, we provide preclinical evidence for the potential use of ECS as an efficacious treatment for alcohol use disorder.

## 1. Introduction

Alcohol is by far the most consumed substance among teens and young adults, as yearly reported by the European Monitoring Centre for Drugs and Drug Addiction (EMCDDA) and the US National Center for Drug Abuse Statistics (NCDAS). Alcohol initiation typically starts at younger ages, during early adolescence, and at higher rates than other drugs of abuse. This is extremely worrisome since the behavioral abnormalities – anxiety, disinhibition, impulsivity, risk-taking, diminished cognitive plasticity – and neural effects of adolescent alcohol use may persist into adulthood (reviewed by Spear, 2016, 2018; Lees et al., 2020), promoting an increased propensity to alcohol use disorder (Strong et al., 2010; Sherrill et al., 2011). This specific vulnerability can be modeled at the preclinical level by administering in early adolescence a binge regimen of ethanol exposure capable of later inducing an increased voluntary ethanol consumption in adult rats (Colom-Rocha et al., 2024), as measured in a two-bottle choice paradigm with unlimited access to a 20% alcohol solution in drinking bottles for 3 consecutive days a week (Colom-Rocha et al., 2023). In this scenario, we could utilize this model of increased propensity to alcohol use disorder to evaluate novel therapeutic approaches.

The treatment of alcohol use disorder is based on several approaches, both non-pharmacological (i.e., evidence-based behavioral health treatments: cognitive behavioral therapy, motivational enhancement therapy, etc.) or pharmacological through approved medications (e.g., disulfiram, acamprosate, naltrexone; reviewed by Koob, 2024). However, since treatment is greatly underutilized due to many challenging factors (reviewed by Koob, 2024), and relapse rates are high, there is a need for providing novel rapid solutions for the treatment of alcohol use disorder (e.g., Gordon et al., 2024). In this context, the use of neuromodulation techniques is becoming more studied for the treatment of substance use disorders (e.g., Mehta et al., 2024; Oesterle et al., 2025), since they aim at stimulating dysfunctional brain circuits (e.g., components of the meso-cortico-limbic pathway) to counterbalance imbalances and reduce substance consumption and craving (e.g., Balconi et al., 2015; Maiti et al., 2017; Steele, 2021).

As for the particular use of neuromodulator approaches for alcohol-use disorder (as opposed to other drugs) most of the interest has been centered in non-invasive neuromodulator approaches (reviewed by Herremans and Baeken, 2012; Trojak et al., 2018; Azevedo and Mammis, 2018; Maatoug et al., 2021; Rosenthal et al., 2024; Oesterle et al., 2025), such as repetitive transcranial magnetic stimulation (rTMS) with proven efficacy reducing craving in alcohol use disorder (e.g., Sorkhou et al., 2022) or transcranial direct current stimulation (e.g., Chmiel and Kurpas, 2025). However, there is a gap in the literature regarding the hypothetical benefits of electroconvulsive therapy for alcohol use disorder. Given our extensive experience inducing electroconvulsive seizures (ECS) for therapeutic purposes in rodent models of psychopathology (García-Fuster and García-Sevilla, 2016; García-Cabrerizo et al., 2020; Ledesma-Corvi and García-Fuster, 2023a,b,c), the present study aimed at evaluating the role of ECS as a therapeutic option for reducing the increased voluntary ethanol consumption caused by early drug initiation in our validated preclinical animal model.

Moreover, in line with our previous study (Colom-Rocha and García-Fuster, 2025), we will evaluate changes in morphological and structural hippocampal markers associated with ethanol exposure and neurotoxicity that might contribute to the development of addictive-like responses and/or treatment response. Particularly, NeuroD will be used as a marker of early neural progenitors during hippocampal neurogenesis (e.g., Nwachukwu et al., 2022; Buján et al., 2024; Colom-Rocha and García-Fuster, 2025); BDNF as a trophic neuroplastic marker (e.g., Peregud et al., 2025; Xiong et al., 2026); neurofilament (NF-L, for light chain) as marker of neuronal shape organization and function (e.g., Beitner-Johnson et al., 1992; Reis et al., 2015; Karoly et al., 2021; Colom-Rocha and García-Fuster, 2025); and finally the ratio between NF-L and BDNF as an index of neurotoxicity (Requena-Ocaña et al., 2023; Colom-Rocha and García-Fuster, 2025).

## 2. Experimental procedures

### 2.1. Animals

For this experiment we used a total of 153 Sprague-Dawley rats (77 males, 76 females) that were bred in our animal facility at the University of the Balearic Islands. After weaning on PND 21, rats were housed in standard cages in groups of 2-4 in a climate-controlled room (22 °C, 70% humidity) with limitless access to a standard diet and water and in a 12 h light/dark schedule (lights on at 8:00 AM). Procedures complied with ARRIVE Guidelines (Percie du Sert et al., 2020), EU Directive 2010/63/EU, and Spanish Royal Decree 53/2013, requiring prior approval by the Local Bioethical Committee (CEEA 238/07/24) and Regional Government (SSBA 19/2024 AEXP). All efforts were made to minimize the number of rats utilized, the number of procedures, which were performed during the light period, and their suffering. In this context, we did not monitor the stages of the estrous cycle, to avoid unnecessary stress only in females. Also, since similar individual variability was observed for males and females in the present study and in line with prior studies (Becker et al., 2016; Kaluve et al., 2022) and considering that cyclicity of females was not part of our research question (e.g., Beltz et al., 2019).

### 2.2. Ethanol exposure during adolescence

Adolescence is a brain developmental period, highly conserved and with similar stages across species; both human (Backes and Bonnie, 2019; Christie and Viner, 2005) and rodent (Spear, 2004) adolescence is divided in early (10 to 13 years vs. postnatal day, PND 21-34), mid (14 to 17 years vs. PND 34-46), and late (18 to 21 years vs. PND 46-59) adolescence respectively. To mimic an early start during the first stages of adolescence, male and female rats were treated in a binge manner with ethanol (2 g/kg, i.p.; 3 rounds of 2 days at 48-h intervals; PND 29-30, PND 33-34 and PND 37-38) or saline (0.9% NaCl, i.p., 1 ml/kg; Fig. 1). This binge administration pattern was based on prior studies from other groups (Pascual et al., 2007, 2017; Crabbe et al., 2011) as well as ours (Colom-Rocha et al., 2024; Colom-Rocha and García-Fuster, 2025). This procedure rendered 4 experimental groups: male-saline (n = 38), male-ethanol (n = 39), female-saline (n = 37) and female-ethanol (n = 39). Note that experimental treatment groups were also combined by sex to provide a broader analysis in terms of future extrapolations to the results to the general population (i.e., mixed-sex cohort). After the adolescent treatment, rats were left undisturbed during forced withdrawal until adulthood (PND 39-78), when they were single housed (PND 79) to later assess voluntary ethanol intake (see Fig. 1).

**Fig. 1.**
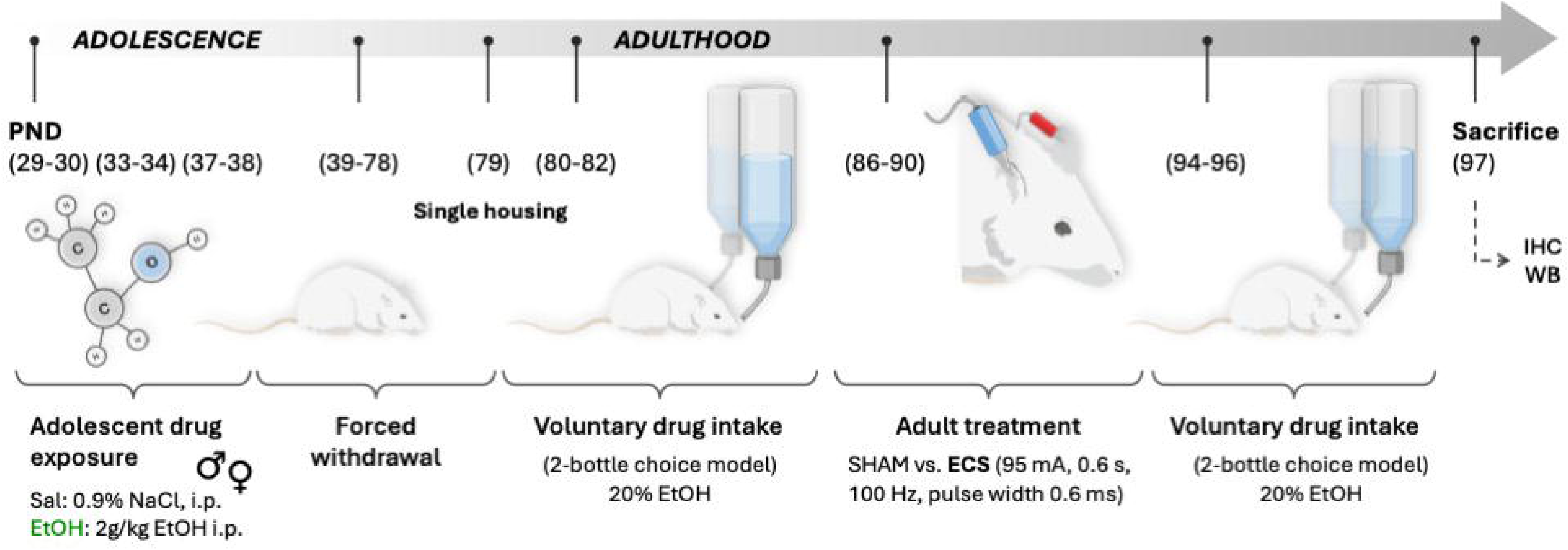
Experimental timeline. Voluntary ethanol consumption in adult rats of both sexes following adolescent drug exposure and/or electroconvulsive seizures (ECS) treatment. Experimental procedures during adolescence (post-natal, PND, 29-38): saline (Sal: 0.9% NaCl, i.p.) or ethanol (EtOH: 2 g/kg, i.p.) during 3 rounds of 2 days at 48-h interval (PND 29-30, 33-34, and 37-38). Experimental procedures during adulthood: single housing rats (PND 79); voluntary consumption of 20% EtOH in a two-bottle preference test over 3 consecutive days (PND 80-82); ECS (95 mA, 0.6 s, 100 Hz, pulse width 0.6 ms) vs. SHAM-treatment during 5 consecutive days (PND 86-90); voluntary consumption of 20% EtOH in a two-bottle preference test over 3 consecutive days (PND 94-96); sacrifice and brain samples collection (PND 97).

### 2.3. Voluntary ethanol consumption following forced withdrawal

Rats exposed to ethanol or saline during adolescence were then allowed to voluntarily drink ethanol (20%) by a two-bottle choice test, for 3 consecutive days (PND 80-82; Fig. 1). This procedure, which provided access to 20% ethanol, was previously described to exert a steady consumption that was sustained over time when the pattern was repeated intermittently week after week (Colom-Rocha et al., 2023). Moreover, prior studies from our group proved that adolescent ethanol exposure (the same binge paradigm) increased voluntary ethanol consumption using this procedure (e.g., Colom-Rocha et al., 2024), and we therefore aimed at replicating these results so we could then ascertain the therapeutic response of ECS. To do so, ethanol bottles were daily placed in alternate positions (right or left side) to prevent any side preferences. Bottles were weighed every morning during the 3 days of ethanol access. Results are expressed (average consumption during all 3 sessions) as ethanol preference (%) and dose (g/kg/24 h). Ethanol preference was calculated as the amount of ethanol consumed (ml) divided by total fluid intake (sum of water + ethanol bottles in ml) and multiplied by 100 (% values).

### 2.4. Neuromodulation treatment through the induction of electroconvulsive seizures (ECS)

Followed by a 4-day period without access to ethanol (early withdrawal), rats were allocated in groups to either receive for 5 consecutive days (PND 86-90), a daily session of ECS (95 mA for 0.6 s at a frequency of 100 Hz square wave pulses, pulse width 0.6 ms; García-Fuster and García-Sevilla, 2016) through ear clip electrodes using a pulse generator (ECT Unit 7801, Ugo Basile, Italy) or SHAM treatment (ear clips placed but no electrical stimulation applied) (Fig. 1). Experimental groups: male-saline-SHAM (n = 21), male-saline-ECS (n = 11), male-ethanol-SHAM (n = 26), male-ethanol-ECS (n = 11), female-saline-SHAM (n = 21), female-saline-ECS (n = 12), female-ethanol-SHAM (n = 25) and female-ethanol-ECS (n = 11). Note that experimental treatment groups were also combined by sex for a broader analysis in terms of future extrapolations to the results to the general population (i.e., mixed-sex cohort). The parameters of ECS have been extensively characterized in adolescent and adult rats of both sexes and are known to reliably reproduce the characteristic tonic and clonic convulsions while inducing antidepressant- and/or neuroplastic-like effects (e.g., García-Cabrerizo et al., 2020; Ledesma-Corvi and García-Fuster, 2023a, 2023b, 2023c).

### 2.5. Voluntary ethanol consumption following ECS treatment

The next week, rats were again allowed to voluntarily drink ethanol (20%) for a total of 3 consecutive days (PND 94-96) through the two-bottle choice test (Fig. 1). The goal was to evaluate the effect of ECS vs. SHAM treatments on the increased voluntary ethanol consumption induced by adolescent drug exposure.

### 2.6. Hippocampal sample collection for neurochemistry evaluations

Brains were collected following rapid decapitation on PND 97, 24 h after removing the access to voluntary ethanol consumption (Fig. 1) in randomly allocated rats from all experimental groups: male-saline-SHAM (n = 9), male-saline-ECS (n = 8), male-ethanol-SHAM (n = 9), male-ethanol-ECS (n = 7), female-saline-SHAM (n = 9), female-saline-ECS (n = 10), female-ethanol-SHAM (n = 9) and female-ethanol-ECS (n = 8). Note that experimental treatment groups were also combined by sex for a broader analysis in terms of future extrapolations to the results to the general population (i.e., mixed-sex cohort). From each collected brain, the left hemisphere was quickly frozen in isopentane at -30 °C to evaluate neural progenitors by immunohistochemistry, while the right hippocampus was freshly dissected and fast-frozen in liquid nitrogen to ascertain neurotoxicity markers by western blot as detailed below.

#### Neural progenitors through immunohistochemistry

NeuroD was used as a marker of neural progenitors which was assessed by immunohistochemistry in the hippocampus (8 sections/slide, 24 sections/animal; Ledesma-Corvi and García-Fuster, 2023c; Colom-Rocha and García-Fuster, 2025). Briefly, we followed standardized procedures in our research group that consisted in postfixing tissue in 4% paraformaldehyde (Merck, Darmstadt, Germany, cat #76240), epitope retrieval in 10% sodium citrate dihydrate (Thermo Fisher Scientific, Waltham, MA, USA, cat #BP327-1) pH 6.0 at 90 °C for 1 h, followed by incubation in 0.3% peroxidase solution (Thermo Fisher Scientific, cat #426000010) and BSA blocking (Merck, cat #A7906), before an overnight incubation with goat anti-NeuroD (1:25000; Santa Cruz Biotechnology, CA, USA, cat #sc-1084) (Ledesma-Corvi and García-Fuster, 2023c; Colom-Rocha and García-Fuster, 2025). The next day, sections were incubated with biotinylated anti-goat secondary antibody (1:1000 respectively, Vector Laboratories, CA, USA, cat #BA-5000), Avidin/Biotin complex (Vectastain Elite ABC kit; Vector Laboratories, cat #PK-6100) and 3,3’-diaminobenzidine (DAB; Merck, cat #D8001) with nickel chloride (Merck, cat #339350) for signal detection. Tissue was dehydrated in graded alcohols, immersed in xylene (Sharlab, Barcelona, Spain, cat #XI0052) and cover-slipped with Permount® (Thermo Fisher Scientific, cat #SP15-500). Once slides were dry and coded, NeuroD +cells were counted in the dentate gyrus region of hippocampus with a Leica DMR light microscope (63x objective lens, 10x ocular lens; 630x magnification). The overall number of NeuroD +cells for each animal was divided by the hippocampal area quantified (mm^2^, as measured with Motic® Images Plus 3.1 software), yielding a value of NeuroD +cells/mm^2^ as previously described (Colom-Rocha and García-Fuster, 2025).

#### Neurotoxicity markers by western blot

As previously described (Colom-Rocha and García-Fuster, 2025), the right part of the hippocampus was prepared as total homogenates (40 μg) that were run by electrophoresis through 10-14% SDS-PAGE mini-gels (Bio-Rad Laboratories, CA, USA), transferred to nitrocellulose membranes, and incubated (overnight, 4 °C) with specific primary antibodies: (1) anti-NF-L (N5139) (1:1000; cat #5139NR4; Sigma-Aldrich, MO, USA); (2) anti-BDNF (1:10000; cat #ab108319; Abcam, Cambridge, UK) and (3) anti-ß-actin (clone AC-15) (1:10000; Sigma-Aldrich, MO, USA). Then, membranes were exposed to specific secondary antibodies linked to horseradish peroxidase (1:5000 dilution; Cell Signaling), ECL chemicals (Amersham, Buckinghamshire, UK), and developed with an autoradiographic film (Amersham ECL Hyperfilm). Films were quantified through densitometric scanning (GS-800 Imaging Calibrated Densitometer, Bio-Rad) and each sample was run at least 3 times in different gels (each gel contained different brain samples from control and treated rats from both biological sexes). Percent changes in immunoreactivity were estimated with respect to control samples in the various gels (100%), and the mean value was used as a final estimate. The ratio between NF-L and BDNF was calculated and used as an index of neurotoxicity (see Colom-Rocha and García-Fuster, 2025). Membranes were reprobed for ß-actin, which was used as a loading control.

### 2.7. Statistical analysis

The software GraphPad Prism (Version 10; GraphPad Software, CA, USA) was used for data analysis and figure plotting. Each figure displays individual rat symbols as well as bars with the mean value ± standard error of the mean (SEM) for each treatment group. The first analysis of data included sex as a biological variable through two- (independent variables: sex and adolescent drug exposure; Supplementary Table S1) or three-way ANOVAs (independent variables: sex, adolescent drug exposure and adult treatment; Supplementary Table S2 and S3). Moreover, a combined analysis considering rats of both sexes as a mixed-sex cohort group was done through unpaired two-tail Student’s *t*-tests or two-way ANOVAs (independent variables: adolescent drug exposure and adult treatment, Supplementary Table S1-S3) as appropriate. The aim was to be able to understand the impact of adolescent drug exposure and/or adult treatment response independently of biological sex, thus allowing for broader conclusions to the general populations even considering potential baseline differences in responses driven by sex. Pair-wise *post-hoc* comparisons were performed when suitable. The level of significance was set at *p* ≤C0.05. Data supporting the present findings will be available upon reasonable request to the corresponding author.

## 3. Results

### 3.1. Exposure to ethanol during adolescence increases voluntary ethanol consumption in adult rats of both sexes without distinction

Adolescent ethanol exposure induced long-term changes in voluntary ethanol consumption in adult rats (Fig. 2), both at the level of ethanol preference (Fig. 2A) and ethanol dose (g/kg/24 h; Fig. 2B), as detailed in Supplementary Table S1, and measured during 3 consecutive days in a week. Although significant differences were observed by sex (Supplementary Table S1), with females showing higher overall ethanol preference (Fig. 2A) and doses of exposure (Fig. 2B), no significant sex x adolescent drug exposure interactions were observed (Supplementary Table S1). Therefore, adolescent ethanol exposure increased voluntary ethanol consumption in adult rats of both sexes without distinction, which was represented by combining rats of both sexes in the right panels of Fig. 2. Particularly, adolescent ethanol exposure increased ethanol preference (+6.7 ± 1.9%, \*\*\**p* < 0.001; Fig. 2A) and the dose of ethanol consumed (+1.3 ± 0.6 g/kg/24 h, \**p* = 0.023; Figure 2B) in adulthood.

**Fig. 2.**
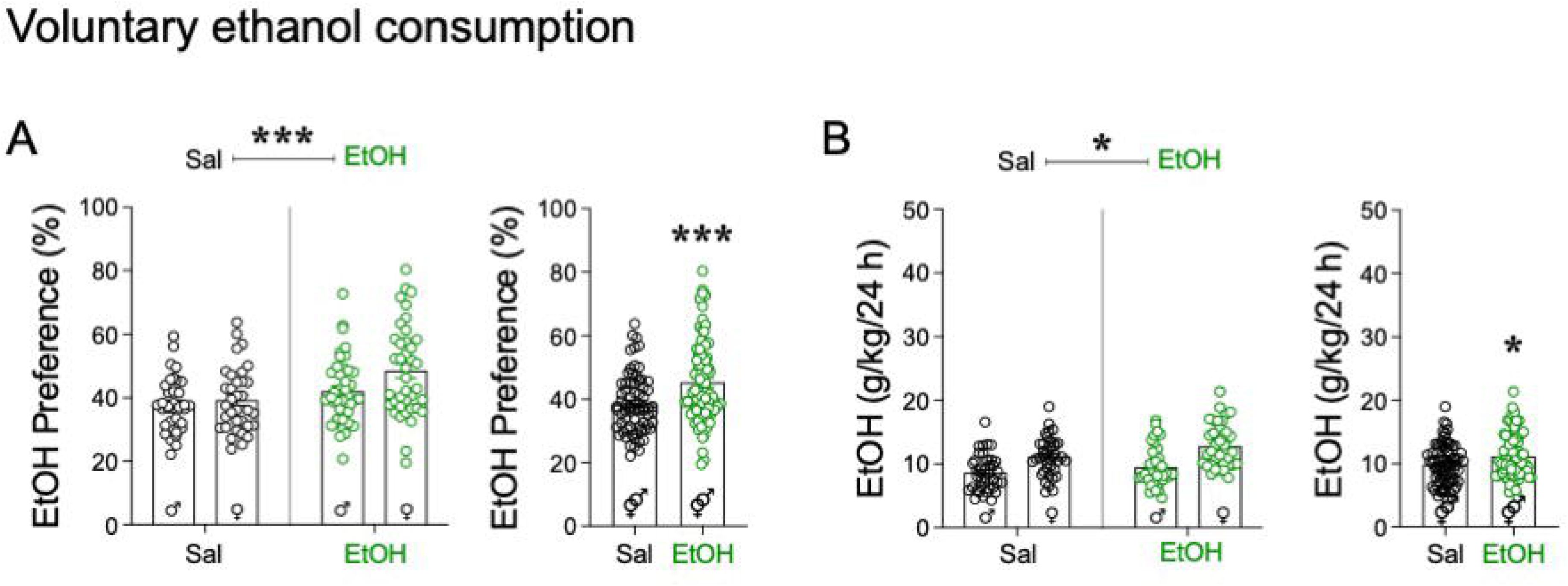
Voluntary consumption of 20% EtOH in a two-bottle preference test over 3 consecutive days. **A** EtOH preference (in %). **B** Daily ethanol dose consumed (g/kg/24h). Groups of treatment: male-saline (n = 38), male-ethanol (n = 39), female-saline (n = 37) and female-ethanol (n = 39). Data represent the mean ± SEM of the preference for ethanol (in %) and the ethanol dose consumed (g/kg/24 h). Each round symbol shows the individual rats (white for saline groups; green for the EtOH group). The left panels included sex as a biological variable (statistical analysis done through two-way ANOVAs), while the right panels evaluated the effect of adolescent drug exposure in a mixed-sex cohort of rats (Student *t*-tests). For further statistical analysis, see Supplementary Table S1. \*\*\**p* < 0.001 and \**p* < 0.05 for representing the overall effects of adolescent ethanol drug exposure and/or when comparing adolescent ethanol exposure vs. saline in a mixed-sex cohort of rats.

### 3.2. ECS decreases voluntary ethanol consumption in adult rats with a prior history of adolescent ethanol exposure

Following ECS vs. SHAM adult treatment, and 2 weeks after their first exposure to voluntary access to ethanol, rats were re-exposed to the 2-bottle choice test. When evaluating ethanol preference, the results showed a significant effect of adolescent drug exposure, no overall effect of sex or adult treatment, but a significant sex x adolescent drug exposure x adult treatment interaction (see Supplementary Table S2; Fig. 3A). Interestingly, Šídák’s multiple comparisons test revealed that adolescent ethanol exposure increased ethanol preference in female-ethanol-SHAM-treated rats (+16.3 ± 13.7%, γγγ*p* < 0.001 vs. female-saline-SHAM-treated rats; Fig. 3A), and that this negative impact was reverted when female rats were treated the previous week with ECS (−13.2 ± 4.5%, ψ*p* = 0.043 female-ethanol-SHAM-vs. female-ethanol-ECS-treated rats; Fig. 3A). No significant pair-wise comparisons were detected for male rats. However, when combining rats of both sexes for analysis, a two-way ANOVA showed an overall increase in ethanol preference caused by prior adolescent ethanol exposure (see Supplementary Table S2), observed in SHAM-treated rats (+8.3 ± 2.7%, \**p* = 0.012; Fig. 3B). However, no significant effect was induced by adult treatment at the level of ethanol preference in adulthood (see Supplementary Table S2; Fig. 3B).

**Fig. 3.**
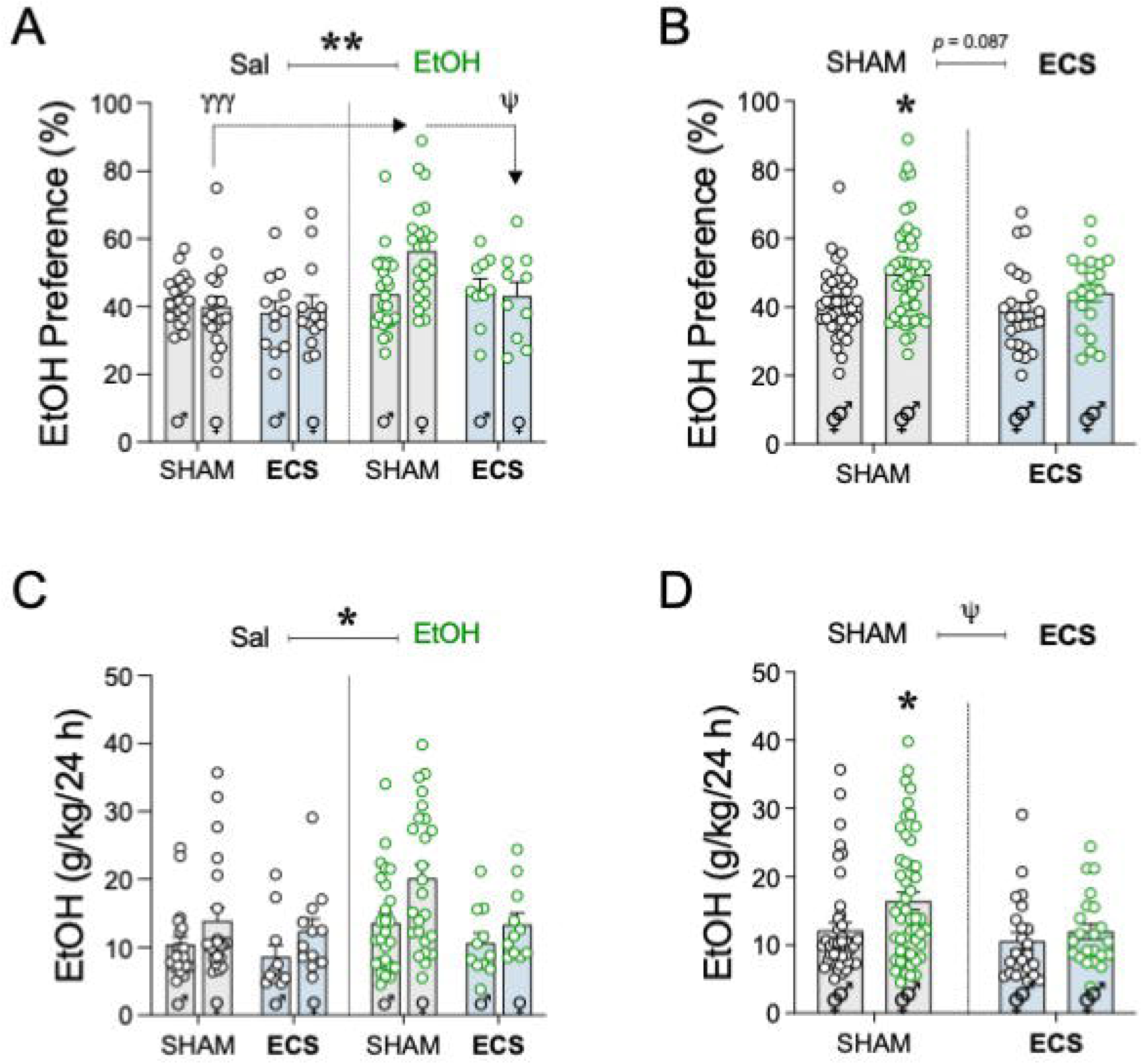
Voluntary consumption of 20% EtOH in a two-bottle preference test over 3 consecutive days following SHAM or ECS treatment. A-B. EtOH preference (in %). **C-D** Daily ethanol dose consumed (g/kg/24h). Groups of treatment: male-saline-SHAM (n = 21), male-saline-ECS (n = 11), male-ethanol-SHAM (n = 26), male-ethanol-ECS (n = 11), female-saline-SHAM (n = 21), female-saline-ECS (n = 12), female-ethanol-SHAM (n = 25) and female-ethanol-ECS (n = 11). Data represent the mean ± SEM of the preference for ethanol (in %) and the ethanol dose consumed (g/kg/24 h). Each round symbol shows the individual rats (white for saline groups; green for the EtOH group) in SHAM- (white bars) vs ECS-(blue bars) treated rats. **A and C** panels included sex as a biological variable (statistical analysis done through three-way ANOVAs), while the **B and D** panels evaluated the effect of adolescent drug exposure and/or adult treatment in a mixed-sex cohort of rats (two-way ANOVAs). For further statistical analysis, see Supplementary Table S2. \*\**p* < 0.01 and \**p* < 0.05 for representing the overall effects of adolescent ethanol drug exposure and/or when comparing adolescent ethanol exposure vs. saline in a mixed-sex cohort of rats. ψ*p* < 0.05 for representing the overall effects of adult treatment.

Interestingly, when evaluating ethanol dose (g/kg/24 h) the results showed significant effects of sex (i.e., females showed higher ethanol doses than males), adolescent drug exposure (i.e., adolescent ethanol drug exposure increased ethanol consumption in adulthood vs. adolescent rats treated with vehicle) and adult treatment (i.e., ECS-showed lower ethanol consumption than SHAM-treated rats; Fig. 3C-D; Supplementary Table S2). When combining rats of both sexes for analysis, a two-way ANOVA showed an overall increase in ethanol preference caused by prior adolescent ethanol exposure (see Supplementary Table S2), observed in SHAM- (+4.3 ± 1.6%, \**p* = 0.041), but not in ECS-treated rats (Fig. 3D). Moreover, there was a significant effect of adult treatment (ψ*p* = 0.032 SHAM-vs. ECS-treated rats; Fig. 3D).

### 3.3. ECS increases hippocampal neural progenitors in adult rats independently of sex and adolescent drug exposure

Brains were evaluated on PND 97, following a first exposure to voluntary ethanol access, ECS vs. SHAM adult treatment, and another exposure to voluntary access to ethanol. When evaluating by immunohistochemistry how these procedures affected the survival of neural progenitors (NeuroD) through a three-way ANOVA, the results showed a significant effect of adult treatment, but no overall effects of sex or adolescent drug exposure (see Supplementary Table S3; Fig. 4A). When the analysis was done combining rats of both sexes, a two-way ANOVA showed the overall increase in hippocampal neural progenitors induced by ECS independently of the adolescent treatment (+313 ± 14 NeuroD +cells/mm^2^, ψψψ*p* < 0.001 SHAM-treated rats; Fig. 4B). Some representative images are shown in Fig. 4C.

**Fig. 4.**
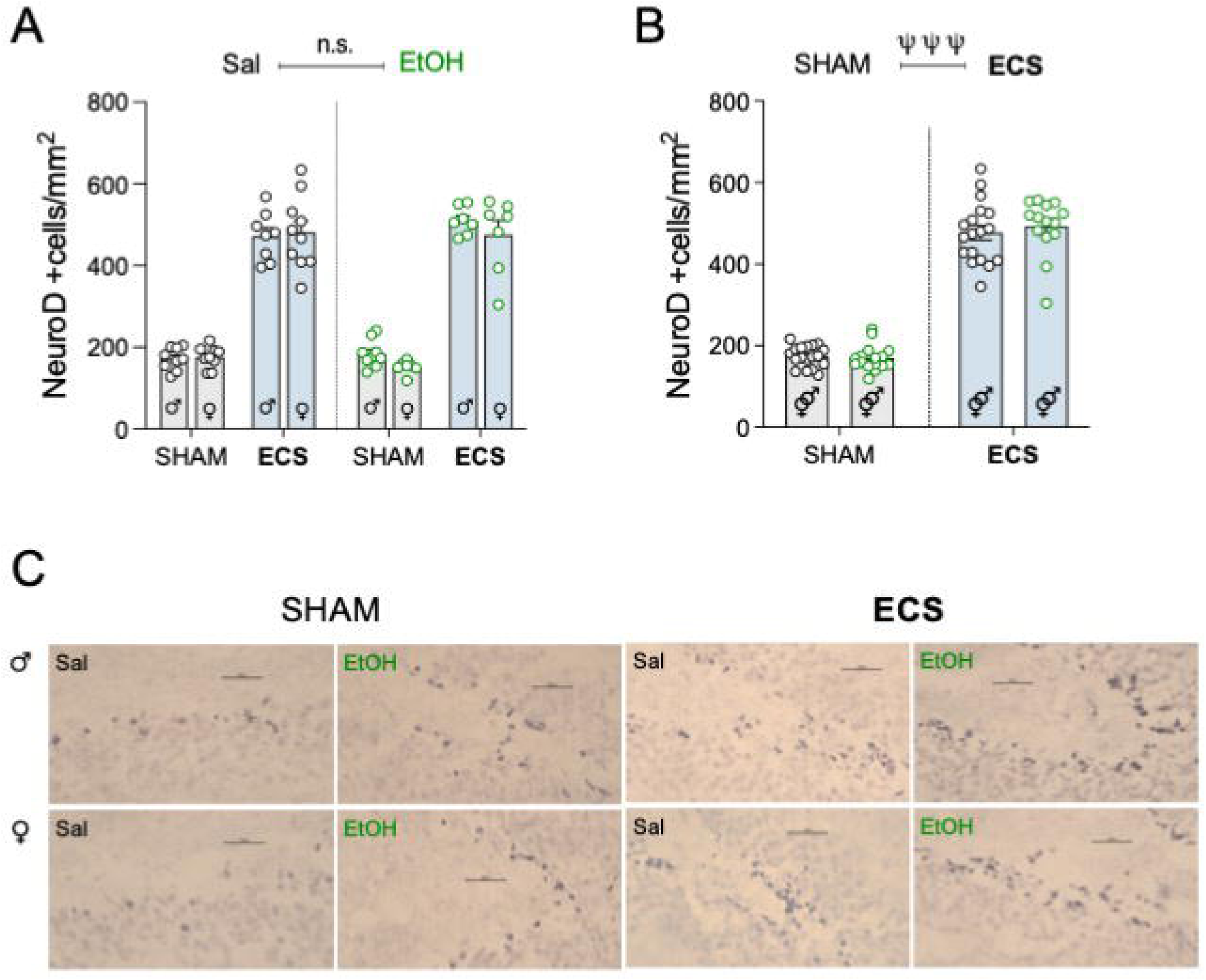
Hippocampal neural progenitors by immunohistochemistry of NeuroD marker. Groups of treatment: male-saline-SHAM (n = 9), male-saline-ECS (n = 8), male-ethanol-SHAM (n = 9), male-ethanol-ECS (n = 7), female-saline-SHAM (n = 9), female-saline-ECS (n = 10), female-ethanol-SHAM (n = 7; 2 rats were excluded from the analysis for methodological problems) and female-ethanol-ECS (n = 7; 1 rat was excluded from the analysis for methodological problems). Data represent the mean ± SEM of the number of NeuroD +cells/mm^2^. Each round symbol shows the individual rats (white for saline groups; green for the EtOH group) in SHAM- (white bars) vs ECS- (blue bars) treated rats. **A** included sex as a biological variable (statistical analysis done through three-way ANOVAs), while **B** evaluated the effect of adolescent drug exposure and/or adult treatment in a mixed-sex cohort of rats (two-way ANOVAs). For further statistical analysis, see Supplementary Table S3. ψψψ*p* < 0.001 for representing the overall effects of adult treatment. **C** Representative images of NeuroD +cells (dark blue labeling in a lighter blue granular layer background) taken with a light microscope using a 63× objective lens. Scale bar: 30 μm.

### 3.4. ECS improves neurotoxic markers in adult rats of both sexes with a prior history of adolescent exposure

The results from the western blot experiments evaluating hippocampal markers (NF-L, BDNF and NF-L/BDNF ratio) through three-way ANOVAs are shown in Supplementary Table S3. For NF-L marker, the results showed a significant effect of adolescent drug exposure, with lower protein levels in rats with a history of adolescent ethanol exposure, but no overall effects of sex or adult treatment (Supplementary Table S3; Fig. 5A). Similarly, when results were evaluated in a combined mixed-sex cohort of rats, there was a significant effect of adolescent drug exposure, with overall lower NF-L protein levels in rats with a history of adolescent ethanol exposure vs. saline-treated rats during adolescence, but no effect of adult treatment (Supplementary Table S3; Fig. 5B).

**Fig. 5.**
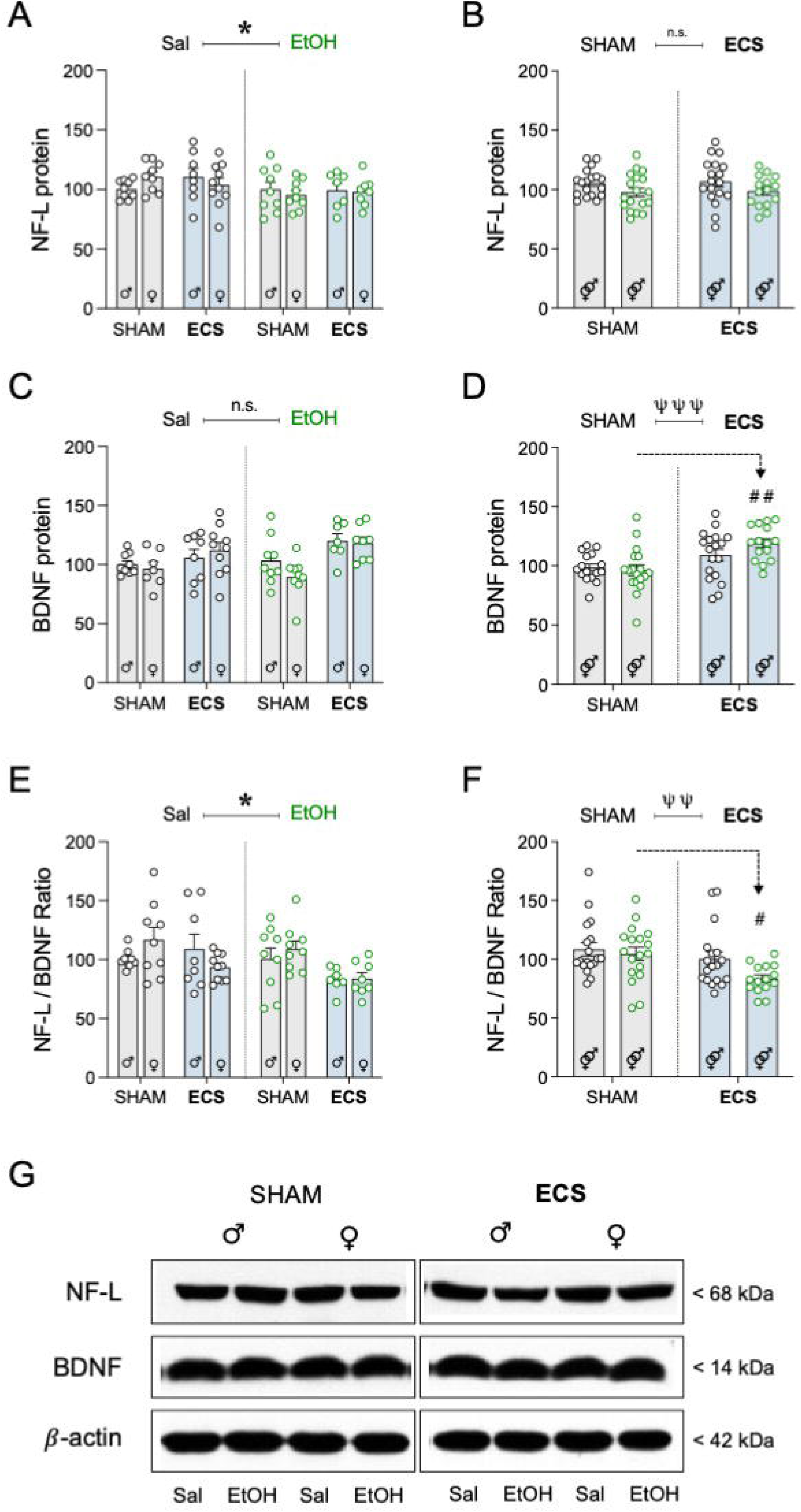
Hippocampal neurotoxicity markers by western blot. Groups of treatment: male-saline-SHAM (n = 9), male-saline-ECS (n = 8), male-ethanol-SHAM (n = 9), male-ethanol-ECS (n = 7), female-saline-SHAM (n = 9), female-saline-ECS (n = 10), female-ethanol-SHAM (n = 9) and female-ethanol-ECS (n = 8). Data represent the mean ± SEM of the corresponding protein under study (**A-B** NF-L, **C-D** BDNF, **E-F** NF-L/BDNF ratio) expressed as % change vs. male-saline-SHAM rats. Each round symbol shows the individual rats (white for saline groups; green for the EtOH group) in SHAM- (white bars) vs ECS-(blue bars) treated rats. **A** included sex as a biological variable (statistical analysis done through three-way ANOVAs), while **B** evaluated the effect of adolescent drug exposure and/or adult treatment in a mixed-sex cohort of rats (two-way ANOVAs). For further statistical analysis, see Supplementary Table S3. \**p* < 0.05 for representing the overall effects of adolescent ethanol drug exposure. ψψ*p* < 0.01 and ψψψ*p* < 0.001 for representing the overall effects of adult treatment. ##*p* < 0.01 and #*p* < 0.05 when comparing ECS-vs. SHAM-treated rats with a history of adolescent ethanol exposure. **G** Representative immunoblots depicting NF-L, BDNF, and β-actin labeling in all experimental groups.

BDNF showed a similar pattern of regulation as NeuroD, since prior studies already showed that ECS increased its levels. Thus, the results showed a significant effect induced by the adult treatment, but no overall effects of sex or adolescent drug exposure (see Supplementary Table S3; Fig. 5C). When the analysis was done combining rats of both sexes, a two-way ANOVA showed no effect of prior adolescent history, while adult treatment increased BDNF (e.g., +22 ± 6.0%, ##*p* = 0.002 when comparing ethanol-SHAM-vs. ethanol-ECS-treated rats; Fig. 5D).

Interestingly, when evaluating the effects over NF-L/BDNF ratio, the statistical analysis by a three-way ANOVA showed significant effects of adolescent drug exposure and adult treatment, but no overall effects of sex (Supplementary Table S3; Fig. 5E). In the mixed-sex cohort of rats (Supplementary Table S3), the effect of prior adolescent history dissipated, while adult treatment still decreased NF-L/BDNF ratio (e.g., -25 ± 7.8%, #*p* = 0.011 ethanol-SHAM-vs. ethanol-ECS-treated rats; Fig. 5F). Representative images for all protein markers are shown in Fig. 5G.

## 4. Discussion

The present study validated our prior data by reassuring that we could reproduce an increased voluntary ethanol consumption in adult rats induced by adolescent ethanol exposure, corroborating that early drug initiation is a risk-factor for later developing an additive-like phenotype. In this scenario we tested the potential beneficial effects of inducing ECS to improve this addictive-prone response. Our results showed that ECS improved ethanol voluntary consumption to levels observed in rats with no prior adolescent ethanol exposure. Moreover, although ECS increased neuroplasticity markers in the hippocampus, such as the number of neural progenitors or BDNF protein content. Interestingly, a prior history of adolescent ethanol exposure impacted NF-L as well as the ratio NF-L/BDNF in adult rats by reducing their hippocampal content as compared to rats treated with saline in adolescence, and ECS treatment further reduced NF-L/BDNF ratio (vs. SHAM-treated rats). Overall, ECS is presented as a great therapeutic option to improve the increased vulnerability to voluntary ethanol consumption in adulthood caused by an early ethanol exposure during adolescence. The results could be generalized for a mixed-sex cohort of rats, allowing for broader conclusions in terms of the potential implications to the general population.

The first part of the study aimed at reproducing a model of addiction-like vulnerability by increasing voluntary ethanol consumption during adulthood caused by early ethanol initiation during adolescence (Colom-Rocha et al., 2024). We previously proved that adolescent ethanol binge exposure was a clear risk-factor for boosting voluntary ethanol preference and consumption in adult rats of both sexes (Colom-Rocha et al., 2024), as measured in a two-bottle choice paradigm with unlimited access to a 20% alcohol solution in drinking bottles for 3 consecutive days a week (Colom-Rocha et al., 2023). Interestingly, in line with prior recent examples showing sex-specific drinking patterns in adult rodents (Foo et al., 2023; McElroy et al., 2023; Colom-Rocha et al., 2023, 2024), female rats were exposed to higher doses of ethanol (g/kg/24 h), since their sizes were considerably smaller than their male counterparts, and although both sexes consumed similar volumes of ethanol (g/24 h). These results aligned with prior data and reinforced the fact that adolescence ethanol initiation could associate with worse addictive-like prognosis in adulthood (Strong et al., 2010; Sherrill et al., 2011; reviewed by Spear, 2016, 2018; Lees et al., 2020). Thus, we have validated and replicated an experimental procedure in which an early adolescent ethanol experience impacted the emerging addictive-like phenotype in adulthood, in line with clinical studies with twins proving that adolescent alcohol use was a risk factor for adult alcohol and drug dependence (Grant et al., 2006). In this validated animal model, we could therefore ascertain potential therapeutic options for preventing alcohol-use disorder caused by early drug initiation.

Against this background, rats with increased voluntary ethanol preference and consumption were either treated with ECS or SHAM following a therapeutic protocol known to induce antidepressant- and/or neuroplastic-like benefits in adult rats (García-Fuster and García-Sevilla, 2016; García-Cabrerizo et al., 2020; Ledesma-Corvi and García-Fuster, 2023c). The results showed that ECS decreases voluntary ethanol consumption in adult rats with a prior history of adolescent ethanol exposure. To the best of our knowledge, this is the first study proving at the preclinical level the benefits of ECS for reducing alcohol consumption in adult rats with a prior history of use in adolescence. Although the results seemed more prominent for female rats, when combining the analysis in a mixed-sex cohort of rats, ECS proved efficacy by reducing ethanol consumption. This is relevant given the need for finding therapeutic options that work broadly for both sexes. The number of studies evaluating neuromodulator approaches for alcohol-use disorder (and/or other substance use disorders) has been increasing with time (reviewed by Herremans and Baeken, 2012; Trojak et al., 2018; Azevedo and Mammis, 2018; Maatoug et al., 2021; Rosenthal et al., 2024; Oesterle et al., 2025), however, most of the interest has been centered to other non-invasive neuromodulator approaches, such rTMS (e.g., Sorkhou et al., 2022), with the hypothetical benefits of ECS being disregarded. Against this background, our novel data postulated ECS as a therapeutic option for reducing alcohol-use disorder caused by early drug initiation as validated in our animal model, suggesting this approach should be further studied to be later implemented into the clinic.

The next part of the study was centered on evaluating hippocampal correlates of ECS response as measured a week post-treatment. The results showed that ECS increased hippocampal neuroplasticity markers (NeuroD and BDNF) in adult rats independently of sex. These results replicated prior studies from our group evaluating the regulation of various stages of hippocampal neurogenesis and BDNF protein content at different times after treatment in rats of both sexes (García-Cabrerizo et al., 2020; Ledesma-Corvi and García-Fuster, 2023c). Interestingly, the beneficial effects of adult ECS treatment on NeuroD were not affected by the prior adolescent drug exposure, even though our prior study proved that adult voluntary ethanol consumption induced reductions in NeuroD (Colom-Rocha and García-Fuster, 2025). The experimental differences in these two studies in terms of the length of exposure to voluntary access to ethanol in adulthood (i.e., 2 weeks vs. 6 consecutive weeks in Colom-Rocha et al., 2024; Colom-Rocha and García-Fuster, 2025) might be responsible for such discrepancies, suggesting that longer weekly access to the drug might be needed in adulthood for the neurotoxic effects to re-emerge following persistent withdrawal after the adolescent exposure (see discussion in Colom-Rocha and García-Fuster, 2025). Remarkably, in rats with a prior history of adolescent ethanol exposure, hippocampal BDNF was significantly increased by ECS as compared to rats treated with SHAM (i.e., ethanol-ECS vs. ethanol-SHAM), suggesting that the beneficial effects of ECS on reducing ethanol’s voluntary intake in adulthood might benefit from the augmentation in BDNF protein content. As for the other neurochemical markers evaluated, adolescent ethanol induced significant long-term reductions in NF-L as compared to rats treated with saline during adolescence, indicative of a persistent neurotoxic impact to the hippocampus emerging following ethanol re-exposure in adulthood. Although ECS did not revert the effect of alcohol on NF-L protein, it significantly decreased the ratio NF-L/BDNF, suggesting a beneficial neuroprotective regulation driven by ECS. This was especially relevant when comparing the results in adult rats with a prior history of ethanol in adolescence (ethanol-SHAM vs. ethanol-ECS). The reduction of NF-L/BDNF might be related to the gain of BDNF trophic action in response to ECS treatment, likely paralleling the observed higher increase of BDNF protein induced by ECS in rats with a history of adolescent ethanol exposure. A reduction of this index could be associated with an improved cognitive impairment (Requena-Ocaña et al., 2023) with potential clinical significance in patients with alcohol use disorders.

In conclusion, reducing adolescence exposure to ethanol is postulated fundamental to avoid future complications related to the development of substance use disorders in adulthood. If this could not be prevented, having access to therapeutic options that could improve the long-term outcome of this early-life experience are of great relevance. In this context, we have provided a novel therapeutic approach based on ECS in a model of increased propensity to alcohol use in adult rats, caused by an early ethanol initiation during adolescence. Moreover, we also suggested a possible role for BDNF in the beneficial effects induced by ECS, especially reducing the neurotoxic ratio NF-L/BDNF. Given the lack of use for ECS in the clinic in the context of substance use disorders, we provide preclinical evidence for an efficacious and safe clinical translation that could have implications for a mixed-sex cohort of subjects, allowing for broader conclusions in terms of the potential implications to the general population.

## Contributors

**Rubén García-Cabrerizo** - Conceptualization; Methodology; Data Curation; Formal Analysis; Writing - Review & Editing.

**Pedro Bergas-Cladera** - Methodology; Formal Analysis; Writing - Review & Editing.

**Carles Colom-Rocha** - Methodology; Writing - Review & Editing.

**M. Julia García-Fuster** - Conceptualization; Data Curation; Formal Analysis; Funding Acquisition; Project Administration; Resources; Original Draft Preparation; Writing - Review & Editing.

All authors contributed to and have approved the final manuscript.

## Supplementary Materials

Supplementary material associated with this article can be found, in the online version.

## References

1. Azevedo CA, Mammis A. Neuromodulation Therapies for Alcohol Addiction: A Literature Review. Neuromodulation. 2018;21:144–148.

2. Backes EP, Bonnie RJ (editors). Adolescent development. The promise of adolescence: realizing opportunity for all youth, National Academies of Sciences, Engineering, and Medicine; Health and Medicine Division; Division of Behavioral and Social Sciences and Education; Board on Children, Youth, and Families; Committee on the Neurobiological and Socio-behavioral Science of Adolescent Development and Its Applications, Washington (DC): National Academies Press (US); 2019 May 16.

3. Balconi M, Finocchiaro R. Decisional impairments in cocaine addiction, reward bias, and cortical oscillation “unbalance”. Neuropsychiatr Dis Treat. 2015;11:777–86.

4. Becker JB, Prendergast BJ, Liang JW. Female rats are not more variable than male rats: a meta-analysis of neuroscience studies. Biol Sex Differ. 2016;7:34.

5. Beitner-Johnson D, Guitart X, Nestler EJ. Neurofilament proteins and the mesolimbic dopamine system: common regulation by chronic morphine and chronic cocaine in the rat ventral tegmental area. J Neurosci. 1992;12:2165–76.

6. Beltz AM, Beery AK, Becker JB. Analysis of sex differences in pre-clinical and clinical data sets. Neuropsychopharmacology. 2019;44:2155–58.

7. Buján GE, D’Alessio L, Serra HA, Guelman LR, Molina SJ. Assessment of hippocampal-related behavioral changes in adolescent rats of both sexes following voluntary intermittent ethanol intake and noise exposure: a putative underlying mechanism and implementation of a non-pharmacological preventive strategy. Neurotox Res. 2024;42:29.

8. Chmiel J, Kurpas D. Neurobiological Mechanisms of Action of Transcranial Direct Current Stimulation (tDCS) in the Treatment of Substance Use Disorders (SUDs)-A Review. J Clin Med. 2025;14:4899.

9. Christie D, Viner R. Adolescent development. BMJ. 2005;330;301–304.

10. Colom-Rocha C, Bis-Humbert C, García-Fuster MJ. Evaluating signs of hippocampal neurotoxicity induced by a revisited paradigm of voluntary ethanol consumption in adult male and female Sprague-Dawley rats. Pharmacol Rep. 2023;75:320–30.

11. Colom-Rocha C, Bis-Humbert C, García-Fuster MJ. Cannabidiol or ketamine for preventing the impact of adolescent early drug initiation on voluntary ethanol consumption in adulthood. Front Pharmacol. 2024;15:1448170.

12. Colom-Rocha C, García-Fuster MJ. Neurotoxic biomarkers of ethanol exposure: from adolescent vulnerability to adult voluntary intake in rats of both sexes. Int J Neuropsychopharmacol. 2025;28:pyaf061.

13. Crabbe JC, Harris RA, Koob GF. Preclinical studies of alcohol binge drinking. Ann N Y Acad Sci. 2011;1216:24–40.

14. Foo JC, Skorodumov I, Spanagel R, Meinhardt MW. Sex- and age-specific effects on the development of addiction and compulsive-like drinking in rats. Biol Sex Differ. 2023;14:44.

15. García-Cabrerizo R, Ledesma-Corvi S, Bis-Humbert C, García-Fuster MJ. Sex differences in the antidepressant-like potential of repeated electroconvulsive seizures in adolescent and adult rats: Regulation of the early stages of hippocampal neurogenesis. Eur Neuropsychopharmacol. 2020;41:132–145.

16. García-Fuster MJ, García-Sevilla JA. Effects of anti-depressant treatments on FADD and p-FADD protein in rat brain cortex: enhanced anti-apoptotic p-FADD/FADD ratio after chronic desipramine and fluoxetine administration. Psychopharmacology (Berl). 2016;233:2955–71.

17. Gordon JA, Volkow ND, Koob GF. No time to lose: the current state of research in rapid-acting psychotherapeutics. Neuropsychopharmacology. 2024;49:10–14.

18. Herremans SC, Baeken C. The current perspective of neuromodulation techniques in the treatment of alcohol addiction: a systematic review. Psychiatr Danub. 2012;24 Suppl 1:S14–20.

19. Kaluve AM, Le JT, Graham BM. Female rodents are not more variable than male rodents: A meta-analysis of preclinical studies of fear and anxiety. Neurosci Biobehav Rev. 2022;143:104962.

20. Karoly HC, Skrzynski CJ, Moe EN, Bryan AD, Hutchison KE. Exploring relationships between alcohol consumption, inflammation, and brain structure in a heavy drinking sample. Alcohol Clin Exp Res. 2021;45 2256–70.

21. Koob GF. Alcohol Use Disorder Treatment: Problems and Solutions. Annu Rev Pharmacol Toxicol. 2024;64:255–275.

22. Ledesma-Corvi S, García-Fuster MJ. Aromatase Inhibition and Electroconvulsive Seizures in Adolescent Rats: Antidepressant and Long-Term Cognitive Sex Differences. Int J Neuropsychopharmacol. 2023a;26:607–615.

23. Ledesma-Corvi S, García-Fuster MJ. Comparing the antidepressant-like effects of electroconvulsive seizures in adolescent and adult female rats: an intensity dose-response study. Biol Sex Differ. 2023c;14:67.

24. Ledesma-Corvi S, García-Fuster MJ. Electroconvulsive seizures regulate various stages of hippocampal cell genesis and mBDNF at different times after treatment in adolescent and adult rats of both sexes. Front Mol Neurosci. 2023c;16:1275783.

25. Lees B, Meredith LR, Kirkland AE, Bryant BE, Squeglia LM. Effect of alcohol use on the adolescent brain and behavior. Pharmacol Biochem Behav. 2020;192:172906.

26. Maatoug R, Bihan K, Duriez P, Podevin P, Silveira-Reis-Brito L, Benyamina A, Valero-Cabré A, Millet B. Non-invasive and invasive brain stimulation in alcohol use disorders: A critical review of selected human evidence and methodological considerations to guide future research. Compr Psychiatry. 2021;109:152257.

27. Maiti R, Mishra BR, Hota D. Effect of high-frequency transcranial magnetic stimulation on craving in substance use disorder: a meta-analysis. J Neuropsychiatry Clin Neurosci. 2017;29:160–71.

28. McElroy BD, Li C, McCloskey NS, Kirby LG. Sex differences in ethanol consumption and drinking despite negative consequences following adolescent social isolation stress in male and female rats. Physiol Behav. 2023;271:114322.

29. Mehta DD, Praecht A, Ward HB, Sanches M, Sorkhou M, Tang VM, Steele VR, Hanlon CA, George TP. A systematic review and meta-analysis of neuromodulation therapies for substance use disorders. Neuropsychopharmacology. 2024;49:649–680.

30. Nwachukwu KN, Healey KL, Swartzwelder HS, Marshall SA. The influence of sex on hippocampal neurogenesis and neurotrophic responses on the persistent effects of adolescent intermittent ethanol exposure into adulthood. Neuroscience. 2022;506:68–79.

31. Oesterle TS, Bormann NL, Al-Soleiti M, Kung S, Singh B, McGinnis MT, Correa da Costa S, Rummans T, Chauhan M, Rojas Cabrera JM, Vettleson-Trutza SA, Scheitler KM, Shin H, Lee KH, Gold MS. Invasive and Non-Invasive Neuromodulation for the Treatment of Substance Use Disorders: A Review of Reviews. Brain Sci. 2025;15:723.

32. Pascual M, Blanco AM, Cauli O, Miñarro J, Guerri C. Intermittent ethanol exposure induces inflammatory brain damage and causes long-term behavioural alterations in adolescent rats. Eur J Neurosci. 2007;25:541–50.

33. Pascual M, Montesinos J, Marcos M, Torres JL, Costa-Alba P, García-García F, Laso FJ, Guerri C. Gender differences in the inflammatory cytokine and chemokine profiles induced by binge ethanol drinking in adolescence. Addict Biol. 2017;22:1829–41.

34. Percie du Sert N, Ahluwalia A, Alam S, Avey MT, Baker M, Browne WJ, et al. Reporting animal research: explanation and elaboration for the ARRIVE guidelines 2.0. PLoS Biol. 2020;18:e3000411.

35. Peregud D, Baronets V, Pavlova O, Pavlov K. BDNF gene polymorphisms and substance use disorders: a systematic review. Rev Neurosci. 2025;37:141–188.

36. Reis KP, Heimfarth L, Pierozan P, et al. High postnatal susceptibility of hippocampal cytoskeleton in response to ethanol exposure during pregnancy and lactation. Alcohol. 2015;49:665–74.

37. Requena-Ocaña N, Araos P, Serrano-Castro PJ, et al. Plasma Concentrations of neurofilament light chain protein and brain-derived neurotrophic factor as consistent biomarkers of cognitive impairment in alcohol use disorder. Int J Mol Sci. 2023;24:1183.

38. Rosenthal A, Haslacher D, Garbusow M, Pangratz L, Apfel B, Soekadar S, Romanczuk-Seiferth N, Beck A. Neuromodulation and mindfulness as therapeutic treatment in detoxified patients with alcohol use disorder. BMC Psychiatry. 2024;24:635.

39. Sherrill LK, Koss WA, Foreman ES, Gulley JM. The effects of pre-pubertal gonadectomy and binge-like ethanol exposure during adolescence on ethanol drinking in adult male and female rats. Behav Brain Res. 2011;216:569–75.

40. Sorkhou M, Stogios N, Sayrafizadeh N, Hahn MK, Agarwal SM, George TP. Non-invasive neuromodulation of dorsolateral prefrontal cortex to reduce craving in alcohol use disorder: A meta-analysis. Drug Alcohol Depend Rep. 2022 Jul 9;4:100076.

41. Spear LP. Adolescent brain development and animal models. Ann NY Acad Sci. 2004;1021:23–6.

42. Spear LP. Consequences of adolescent use of alcohol and other drugs: studies using rodent models. Neurosci Biobehav Rev. 2016;70:228–243.

43. Spear LP. Effects of adolescent alcohol consumption on the brain and behaviour. Nat Rev Neurosci. 2018;19:197–214.

44. Steele VR. A circuit-based approach to treating substance use disorders with noninvasive brain stimulation. Biol Psychiatry. 2021;89:944–6.

45. Strong MN, Yoneyama N, Fretwell AM, Snelling C, Tanchuck MA, Finn DA. Binge drinking experience in adolescent mice shows sex differences and elevated ethanol intake in adulthood. Horm Behav. 2010;58:82–90.

46. Trojak B, Sauvaget A, Fecteau S, Lalanne L, Chauvet-Gelinier JC, Koch S, Bulteau S, Zullino D, Achab S. Outcome of Non-Invasive Brain Stimulation in Substance Use Disorders: A Review of Randomized Sham-Controlled Clinical Trials. J Neuropsychiatry Clin Neurosci. 2017;29:105–118.

47. Xiong JW, Dou MY, Wang Y, Zeng T, Qi X, Wei JN, Shi XW, Cui DD, Dai HZ, Du CY, Xu XM, Wang XF, Zhu X, Guan Y. BDNF restores impaired long-term potentiation of GABAergic synapses induced by chronic ethanol exposure in the VTA and attenuates reward-seeking behavior. Mol Psychiatry. 2026. doi: 10.1038/s41380-026-03532-4.

