## Supplemental Materials for "Electroconvulsive seizures for alcohol use disorder: a preclinical study"

### Supplementary Materials

**Table S1.** Two-way ANOVAs analyses (independent variables: Sex and Adolescent Drug Exposure) including F (DFn, DFd) and *p* values, for data represented in Fig. 2. Color-shadow boxes represent statistically significant comparisons for the variables under study.

Voluntary ethanol consumption (1st week) in adult rats of both sexes

| Behavioral outcomes | | Sex | Adolescent Drug Exposure | Interaction |
|---|---|---|---|---|
| **Fig. 2A** | EtOH Preference (%) | F (1, 145) = 4.49; *p* = 0.036 | F (1, 145) = 13.24; ****p* < 0.001 | F (1, 145) = 1.83; *p* = 0.179 |
| **Fig. 2B** | EtOH (g/kg/24 h) | F (1, 149) = 33.50; *p* < 0.001 | F (1, 149) = 6.26; **p* = 0.014 | F (1, 149) = 0.60; *p* = 0.439 |

**Table S2.** Three-way (independent variables: Sex, Adolescent Drug Exposure and Adult Treatment), or two-way (Adolescent Drug Exposure and Adult Treatment) ANOVAs analyses including F (DFn, DFd) and *p* values, for data represented in Fig. 3. Color-shadow boxes represent statistically significant comparisons for the variables under study.

Voluntary ethanol consumption after treatment in adult rats of both sexes

| Behavioral outcomes | | Sex | Adolescent Drug Exposure | Adult Treatment | Interaction |
|---|---|---|---|---|---|
| **Fig. 3A** | EtOH Preference (%) | F (1, 124) = 1.22; *p* = 0.272 | F (1, 124) = 10.14; ***p* = 0.002 | F (1, 124) = 3.51; *p* = 0.063 | F (1, 124) = 4.61; *p* = 0.034 |
| **Fig. 3C** | EtOH (g/kg/24 h) | F (1, 130) = 9.32; *p* = 0.003 | F (1, 130) = 5.22; **p* = 0.024 | F (1, 130) = 5.78; *p* = 0.018 | F (1, 130) = 0.55; *p* = 0.461 |

Voluntary ethanol consumption after treatment in a mixed-sex cohort of adult rats

| Behavioral outcomes | | Adolescent Drug Exposure | Adult Treatment | Interaction |
|---|---|---|---|---|
| **Fig. 3B** | EtOH Preference (%) | F (1, 128) = 8.72; ***p* = 0.004 | F (1, 128) = 2.97; *p* = 0.087 | F (1, 128) = 0.48; *p* = 0.491 |
| **Fig. 3D** | EtOH (g/kg/24 h) | F (1, 133) = 4.22; **p* = 0.042 | F (1, 133) = 4.71; ψ*p* = 0.032 | F (1, 133) = 1.06; *p* = 0.305 |

**Table S3.** Three-way (independent variables: Sex, Adolescent Drug Exposure and Adult Treatment), or two-way (Adolescent Drug Exposure and Adult Treatment) ANOVAs analyses including F (DFn, DFd) and *p* values, for data represented in Fig. 4 and 5. Color-shadow boxes represent statistically significant comparisons for the variables under study.

Regulation of hippocampal markers after treatment in adult rats of both sexes

| Biomarker | | Sex | Adolescent Drug Exposure | Adult Treatment | Interaction |
|---|---|---|---|---|---|
| **Fig. 4B** | NeuroD +cells/mm2 | F (1, 58) = 0.96; *p* = 0.331 | F (1, 58) = 0.12; p = 0.726 | F (1, 58) = 523.8; *p* < 0.001 | F (1, 58) = 0.02; *p* = 0.893 |
| **Fig. 5B** | NF-L protein | F (1, 61) = 0.02; *p* = 0.897 | F (1, 61) = 4.87; **p* = 0.031 | F (1, 61) = 0.13; *p* = 0.723 | F (1, 61) = 1.90; *p* = 0.173 |
| **Fig. 5D** | BNDF protein | F (1, 59) = 0.65; *p* = 0.647 | F (1, 59) = 0.96; *p* = 0.332 | F (1, 59) = 15.29; *p* < 0.001 | F (1, 59) = 0.01; *p* = 0.927 |
| **Fig. 5F** | NF-L/BDNF protein | F (1, 61) = 0.24; *p* = 0.626 | F (1, 61) = 4.03; ψ*p* = 0.049 | F (1, 61) = 7.13; *p* = 0.010 | F (1, 61) = 1.28; *p* = 0.262 |

Regulation of hippocampal markers after treatment in a mixed-sex cohort of adult rats

| Biomarker | | Adolescent Drug Exposure | Adult Treatment | Adolescent x Adult |
|---|---|---|---|---|
| **Fig. 4B** | NeuroD +cells/mm2 | F (1, 62) = 0.17; *p* = 0.681 | F (1, 62) = 534.2; ψψψ*p* < 0.001 | F (1, 62) = 0.52; *p* = 0.473 |
| **Fig. 5B** | NF-L protein | F (1, 65) = 4.74; *p* = 0.033 | F (1, 65) = 0.09; *p* = 0.763 | F (1, 65) = 0.01; *p* = 0.964 |
| **Fig. 5D** | BNDF protein | F (1, 63) = 0.82; *p* = 0.368 | F (1, 63) = 15.51; ψψψ*p* < 0.001 | F (1, 63) = 2.06; *p* = 0.156 |
| **Fig. 5F** | NF-L/BDNF protein | F (1, 65) = 3.64; *p* = 0.061 | F (1, 65) = 7.43; ψψ*p* = 0.008 | F (1, 65) = 1.47; *p* = 0.231 |
